# Dynamics on the web: spiders use physical rules to solve complex tasks in mate search and competition

**DOI:** 10.1101/2021.01.10.426127

**Authors:** Amir Haluts, Sylvia F. Garza Reyes, Dan Gorbonos, Alex Jordan, Nir S. Gov

## Abstract

A long-standing question in animal behaviour is how organisms solve complex tasks. Here we explore how the dynamics of animal behaviour in the ubiquitous tasks of mate-search and competition can arise from a physics-based model of effective interactions. Male orb-weaving spiders of the genus *Trichonephila* are faced with the daunting challenge of entering the web of a much larger and potentially cannibalistic female, approaching her, and fending off rival males. The interactions that govern the dynamics of males within the confined two-dimensional arena of the female’s web are dominated by seismic vibrations. This unifying modality allows us to describe the spiders as interacting active particles, responding only to effective forces of attraction and repulsion due to the female and rival males. Our model is based on a detailed analysis of experimental spider trajectories, obtained during the approach of males towards females, and amidst their interactions with rival males of different sizes. The dynamics of ’spider particles’ that emerges from our theory allows us to explain a puzzling relationship between the density of males on the web and the reproductive advantages of large males. Our results provide strong evidence that the simple physical rules at the basis of our model can give rise to complex fitness related behaviours in this system.

## Introduction

Intuitively, only sophisticated entities with high cognitive capacity seem capable of solving complex tasks. However, apparently simple organisms find solutions to tasks of striking complexity in various aspects of life, from use of complex acoustic patterns in mate recognition by Tungara frogs[1, 2], to cooperative behaviour of cargo-transporting ants[3, 4], and problem-solving by slime moulds[5, 6]. How relatively simple organisms solve complex tasks is a long-standing question in the study of animal behaviour. Understanding how complex behaviours can arise from simple underlying rules is therefore a primary and enduring goal of behavioural studies.

One example of simple agents solving a complicated–and dangerous–task comes in the mate search behaviour of orbweaving spiders of the genus *Trichonephila*. In these spiders, females build large prey-capture webs, to which mature males migrate in search for a mate[7, 8]. In the confined arena of the female’s web, a male is faced with a challenging task. It has to avoid being eaten by the potentially cannibalistic female, which can be an order of magnitude larger than the male[7, 9, 10], and to compete for mating opportunities with rival males that often arrive at the same web[11, 12]. Attaining reproductive success in this environment is a complex challenge, that mirrors a classic optimal foraging problem[13, 14, 15]. Males who forage for a mate must choose whether to remain on a resource patch – their residence female’s web, or move to a new web, all the while accounting for local and global levels of competition.

Remarkably, when faced with this non-trivial optimization problem, *Trichonephila* males are able to find solutions that achieve higher than expected fitness payoffs[12]. That these males can find such solutions is all the more impressive in the context of cognition and performance in small animals with miniaturised nervous systems[16, 17]. The small male spiders, navigating competition, risk, and reward, nevertheless show sophisticated behaviours that belie their size and apparent neuronal capacity[18, 19]. The difficulties facing these males are exacerbated by their limited sensory capacities. Like most other web-building spiders, male *Trichonephila* have very poor vision, and rely almost exclusively on a single seismic modality – web vibrations – to interact with their environment[20, 21, 22].

Here we ask how these apparently simple spider agents solve the complex challenge that they face on the web. Specifically, we ask whether spiders may use relatively simple strategies that require no cognition or memory, but instead are based only on responses to physical forces. The confined two-dimensional arena of the female’s web is a unique natural setup, where the dynamics of interacting agents can be tracked and studied with great detail. We rely on systematic analysis of male trajectories and behavioural observations in *Trichonephila clavipes* spiders to tease apart the underlying physical rules of male-female and male-male interactions. Based on these rules, which amass to simple responses to vibrational cues and geometric information, we construct a complete physical model for the dynamics of male spiders on a female’s web. The model is framed in terms of effective ’spider potentials’, that give rise to effective interaction forces between the spider agents. Treating the males as interacting active Brownian particles (or ’spider particles’), we insert these effective potentials into Langevin equations that govern their motion in a two-dimensional web-space.

Our theoretical framework gives rise to male agent dynamics that stand in quantitative agreement with the motion of real males. Within the same framework, we introduce an empirically motivated body-size dependence into our model for a male’s interaction potential. We thereby explore the outcomes of differing competitive abilities in male-male contests, and the way in which these outcomes vary with respect to the density of males on the web. Our results shed light upon previous observations on the relation between size advantage and male density, and demonstrate that the solution to a complex ecological task can emerge from a few simple interaction rules that originate from a unifying modality.

## Results

### Male-female interactions and single-male dynamics

Understanding how web architecture and the presence of the female influence the dynamics of single males on the web is the starting point for our theoretical description. During most of the day, the female assumes a static, down-facing position at a dedicated location in the structural center (’hub’) of the orb, and waits for prey to get caught[23, 24]. Females are highly reactive towards vibratory signals that they interpret as potential prey captures, and this attitude is most prominent when the source of vibrations is within the main prey capture zone of the web, which occupies a wide sector in front of the hub-dwelling female[25, 26] (’frontal sector’, Fig.1a). *T. clavipes* males may have evolved to utilize the well-defined and conserved architecture of female webs to navigate their way towards the female and minimize the pre-mating risk of predation.

**Figure 1.**
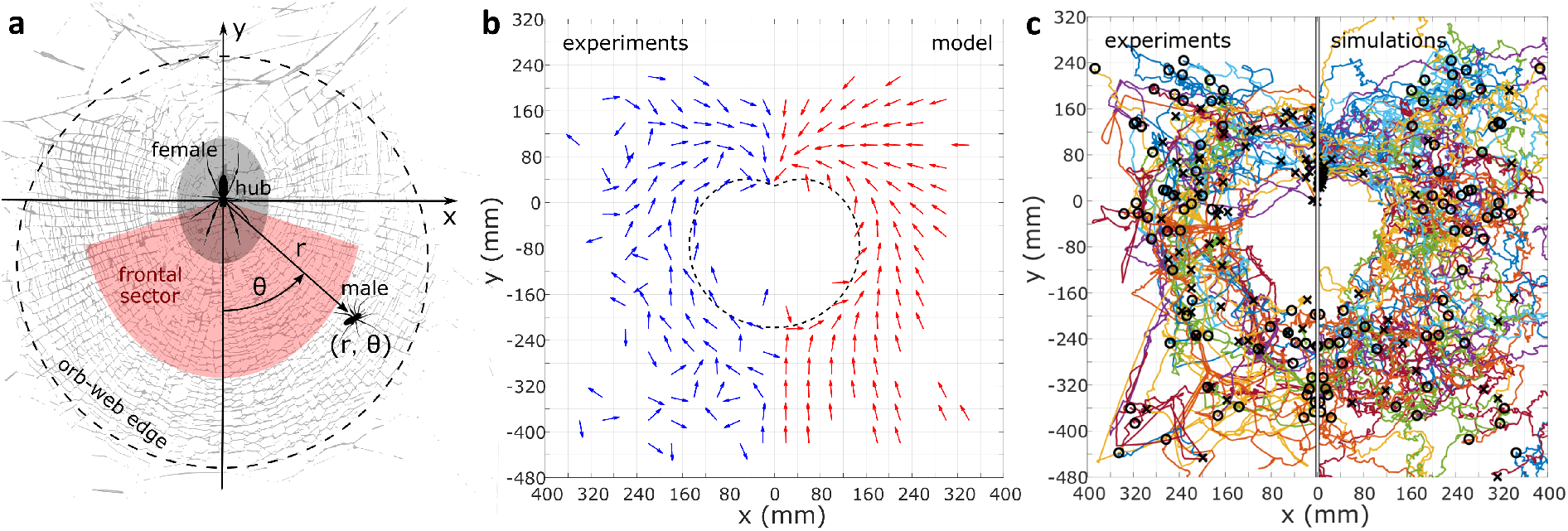
Single-male dynamics. **(a)** Female-centered coordinate system. The head of the static, down-facing female is taken as the origin. The position of a male in this reference frame is shown in polar coordinates. Colored areas roughly illustrate the location and shape of the ’hub’ and the ’frontal sector’. The dashed circle approximates the rim of the orb-web, within which real males can effectively travel continuously. **(b)** Flow fields comparison. The normalized gradient field (director field) of the optimized female potential (Eq. (1) with *β/α* = 94, *C* = 1.32) is shown alongside its experimental analogue (calculated directly from the experimental trajectories of **c**), to allow for a visual comparison. Each director indicates the mean local direction of trajectories that pass through its bin, bordered by the grid square. The dashed contour indicates where the radial partial derivative of the female potential is zero (*∂V*_fem_/*∂r* = 0). **(c)** Single-male trajectories comparison (*N* = 59). Each experimental trajectory was obtained from an independent experiment, in which a single male was placed at some initial position, and left to move freely, in the presence of the female, for about 10 minutes. The simulated single-male trajectories were produced for the same initial positions and durations as the experimental ones. Initial and final positions are marked by circles and x’s, respectively. In **b** and **c**, the data is ’folded’ along the axis of symmetry (the y-axis) onto one half-plane. In **b**, directors in spatially equivalent bins were further averaged post-folding. See Supplementary Fig. 2 for the unfolded experimental trajectories.

Indeed, males exhibit non-trivial dynamics in their motion on the web. In agreement with [25], we observed that males tend to avoid the frontal sector, and approach the female from behind, where it is least reactive. In single-male experiments (Methods and Supplementary Figs. 1,2), we demonstrate that this behaviour leads to effective ordered ’flow’ of males around the female and towards a desired spot within the hub (Fig. 1b,c), where a male would wait for an opportunity to copulate. Note that due to the natural right-left symmetry of the orb system (Supplementary Fig. 2), we could fold the data onto one half-plane, and thereby improved the statistics of this analysis.

Following a mean-field approach, we treat the observed effective flow of males as a global flow-field associated with the influence of the female and its web. In the absence of rival males, this global flow-field governs the motion of a male on the web. Since the position of the female is essentially fixed in the time scales of interest, of the order of minutes to hours, the influence of the female and its web can in turn be thought of as an effective ’female potential’, *V*_fem_. This effective potential describes an averaged effect on approaching males. Since the motion of males on the web is mostly highly overdamped, we neglect inertial effects, and take the local velocity of isolated males to be directly proportional to the local gradient of *V*_fem_.

As the males are evidently attracted to the female, but avoid the frontal sector (Fig. 1b,c), it is natural to construct *V*_fem_ as a combination of two components – isotropic attraction and anisotropic repulsion. The isotropic attraction is associated with the movement of males towards the female from the edges of its web, and is therefore long-range with respect to the dimensions of the system. The anisotropic repulsion expresses the observed avoidance of the frontal sector by approaching males, that are aiming to evade detection by the female. In recent experiments we found that the anisotropic repulsion originates from web architecture, and therefore exists even if the female is absent, while the attraction is directly associated with the presence of the female in the web (Garza et al., in prep.).

The combination of isotropic and anisotropic contributions to the effective potential, that acts in the two-dimensional (2D) surface of the the web, suggests to write *V*_fem_ as the leading terms of a 2D multipole expansion[27]. The leading order long-range (monopole) term corresponds to the isotropic attraction, while the next-order (dipole-like) term corresponds to the anisotropic repulsion (Supplementary Fig. 3a,b). Namely, we propose to describe *V*_fem_ by the potential

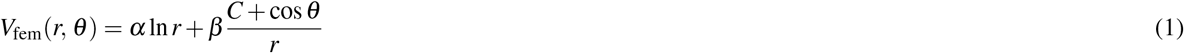

where (*r, θ*) is the position of the male in the female-centered coordinate system, as defined in Fig. 1a, and *α, β* > 0. Note that in the dipole-like term we added a constant *C* ≥ 1 to the numerator. This constant accounts for the fact that the signal represented by this term is never attractive. The potential of Eq. (1) is visually represented in Supplementary Fig. 3c,d.

The effective force derived from Eq. (1), given by −∇*V*_fem_, generates a flow-field with two fixed points – one of which is globally stable, and the other is an unstable saddle (Supplementary Fig. 3c and Supplementary Note 1). These properties of *V*_fem_ can be directly related to their experimental analogues. The globally stable fixed point is associated with the location of the hub, and the unstable saddle point is associated with the boundary of the frontal sector directly in front of the down-facing female (Fig. 1a), as they are perceived by males. By comparing the normalized flow-field (director field) of *V*_fem_, given by −∇*V*_fem_/|∇*V*_fem_|, with its experimental analogue, obtained from single-male trajectories (Fig. 1b), we optimized the parameters of *V*_fem_ such that the directional discrepancy between the two flow-fields is minimal (Supplementary Notes 2,3 and Supplementary Fig. 4). Note that *V*_fem_ has only two independent parameters, *C* and the ratio *β/α*, as can be shown by rescaling (Supplementary Notes 2,3).

Compared to the deterministic flow-field generated by −∇*V*_fem_, experimental single-male trajectories are characterized by positional noise and notable directional persistence, as depicted qualitatively in Fig. 1c To incorporate these characteristics into our theoretical framework, we treat the males as active Brownian particles[28] in an external effective potential, governed by the following Langevin equations[29]

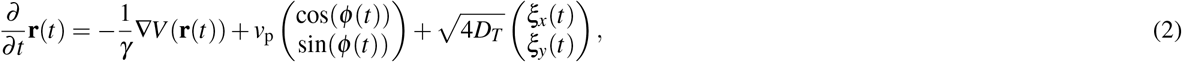

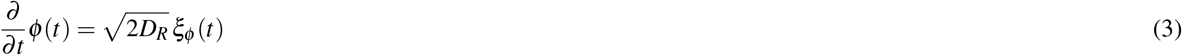

where **r**(*t*) is the position and *ϕ* (*t*) is the persistence angle of a male ’spider particle’. In Eq. (2), −∇*V*(**r**(*t*)) is the deterministic force felt by the male in position **r**(t) due to a potential *V*, and *γ* is an effective friction coefficient for the motion of a male on the web. The second term accounts for the internal directional persistence of the spider, where *v*p is its mean persistence speed. The third term is the stochastic component of the translational motion, where *D_T_* is an effective translational diffusion coefficient, and *ξ_x_*(*t*), *ξ_y_*(*t*) are the stochastic sources for each of the spatial dimensions. In Eq. (3), *D_R_* is an effective rotational diffusion coefficient, and *ξ_ϕ_*(*t*) is the corresponding stochastic source. All the stochastic sources are standard Gaussian white noises, and are mutually uncorrelated.

The four Langevin parameters, *γ, v_p_, D_T_*, and *D_R_*, determine the local characteristics of motion of our model’s ’spider particles’. We estimated the values of these parameters by quantitative analysis of experimental single-male trajectories and comparison to simulated trajectories, generated by integrating Eqs. (2) and (3) with *V* = *V*_fem_ (Supplementary Note 4 and Supplementary Fig. 5). A comparison between experimental and simulated male trajectories is shown in Fig. 1c as a visual validation of our theoretical framework thus far. This validation relies on the already optimized female potential, and therefore establishes the effective physics that governs the motion of single males. Note that our simulations successfully describe the overall shape and spatial distribution of full trajectories, while the temporal dynamics of real males are affected by sporadic pauses, that are not included in the model. We used Eqs. (2) and (3) with a first-order (Euler) integration scheme (as described in [29]) for all simulations in this work.

### Male-male interactions and inter-male dynamics

Female webs serve as confined interaction arenas for competing males, as multiple males often occupy the same web[11], and compete with each other for mating opportunities. By constructing *V*_fem_, we obtained the effective potential surface on which male-male interactions take place. In this section, we describe the physical implications of inter-male competition and extend our theoretical framework to account for inter-male dynamics.

Males display several levels of aggression towards rivals to obtain and retain the desired hub position. Previous behavioural studies have described the escalation in aggression as the distance between the interacting males decreases[8, 25, 30]. When two males first become aware of one another, they employ vibratory threats. In this type of interaction, the males try to intimidate each other by actively jerking the web to generate vibrations[25]. If the initial vibratory threats did not yield the desired response, a persistent aggressor will move directly towards its rival. The rival either retreats, or becomes engaged with the aggressor and prepares for a fight[11]. Finally, if both males persist, they will come into contact and grapple[30], in an effort to forcefully chase their opponent away[8]. We observed that males keep employing vibratory threats as they get closer to each other, probably as means of resolving the competition without the ultimate escalation to grappling.

To understand the dynamics of inter-male competition, we conducted controlled two-male experiments, in which we allowed two males to interact in a female’s web (Methods; Fig. 2a; Supplementary Movie 1). At the beginning of each trial, two males of similar sizes were placed near the rim of the orb, far enough from one another to not be initially engaged in an interaction (Supplementary Fig. 1). In most of these trials, the males reached the threshold inter-male distance for aggressive engagement, mostly as a consequence of their effort to reach the female, and competitive interactions took place (Fig. 2a,b). From these experiments, we extracted the averaged inter-male relative velocity as a function of inter-male distance *d* (’inter-male velocity profile’), which serves as a direct quantitative measure for the physical nature of male-male interactions (Fig. 2c).

**Figure 2.**
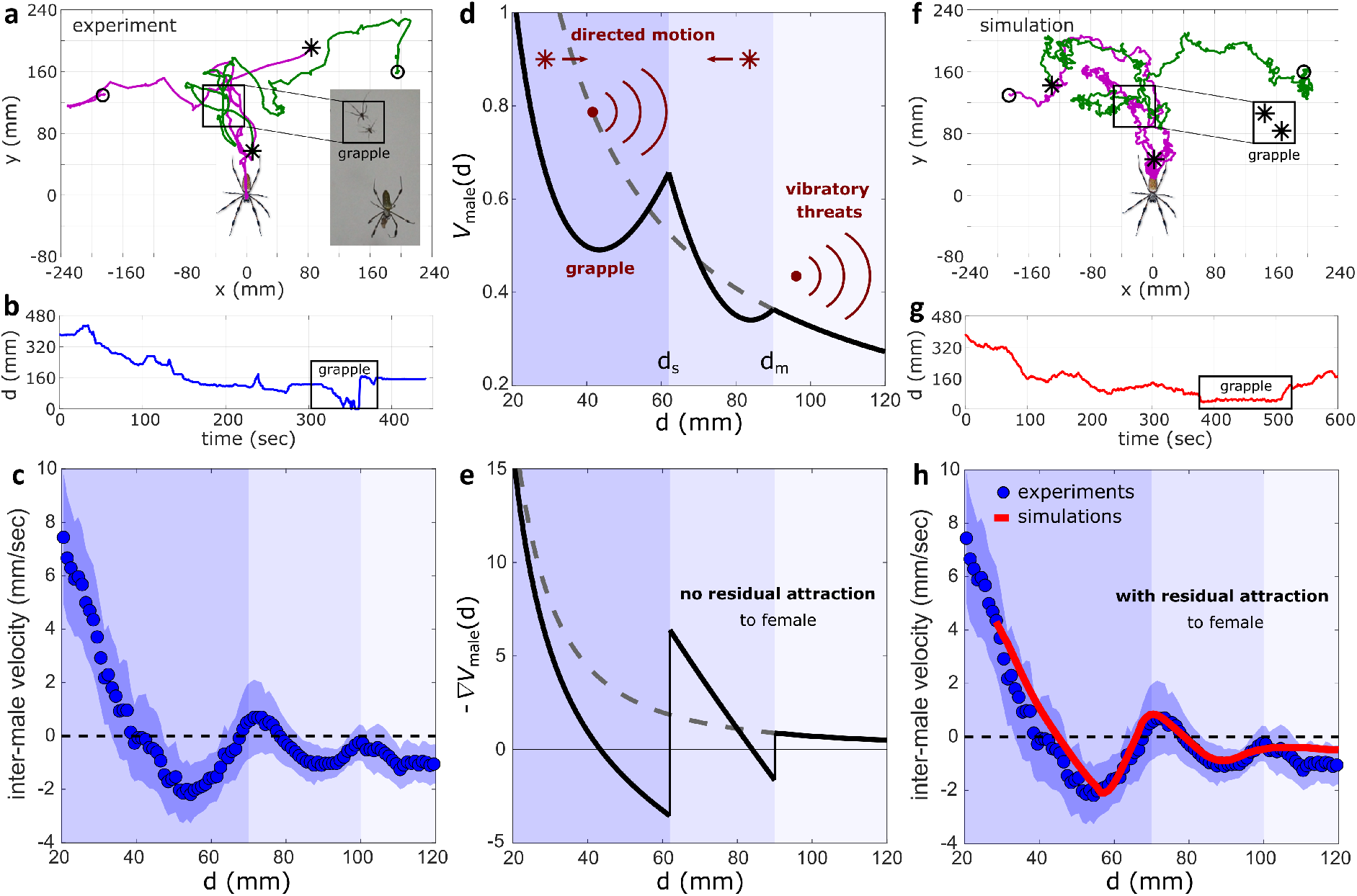
Two-male dynamics. **(a)** Example of male trajectories from a two-male experiment. In these experiments, two males were placed in a female’s web, near the rim of the orb’s upper part, and left to freely move and interact. This particular instance features a grapple between the males (snapshot in inset), which resulted in a swap for the hub position. Initial and final positions are marked by circles and asterisks, respectively. **(b)** The distance between the males in **a** throughout the experiment. **(c)** Averaged inter-male (relative) velocity as a function of inter-male distance *d* (*N* = 29). Silhouette shows SEM. **(d)** The male effective potential *V*_male_ (Eq. (4)), with the parameter values: *δ* = 32.7, *k* = 19, 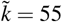, *d*_0_ = 0, 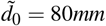, *d*_s_ = 62mm, *d*_m_ = 90mm. **(e)** The force profile derived from *V*_male_. In **d** and **e**, the grey dashed line extrapolates the vibration-based repulsive term (*δ/d* and *δ/d*^2^, respectively). **(f)** An example of male trajectories from a two-male simulation, with initial positions as in **a**. **(g)** The distance between the males in **f** throughout the simulation. **(h)** Inter-male velocity profiles comparison. The red profile was obtained from simulations as in **f** (*N* = 2000), with parameter values as in **d**. Velocities were calculated with a time window of 2*sec* (50 video frames or simulation steps). In **c**-**e** and **h**, the different-shaded bands indicate the 3 distance-dependent regimes of inter-male interaction. The bands in **d,e** correspond to the optimized values of *d*_s_ and *d*_m_, and the bands in **c** and **h** are visual estimates. In **c** and **h**, a symmetric averaging window of length 11mm was used for smoothing.

In agreement with the behavioural observations described above, the inter-male velocity profile reveals three distinct interaction regimes (Fig. 2c). We attribute positive inter-male velocity (a tendency to increase *d*) to effective repulsion between the males, and negative inter-male velocity (a tendency to decrease *d*) to effective attraction, which is interpreted, in accordance with observations, as escalation in aggression. In analyzing Fig. 2c, one should consider the residual attraction of both males to the female, which shifts their relative velocity towards more negative values. This effect is most prominent when the males are far apart and not strongly engaged with each other (as in the right-most region of Fig. 2c), so that the attraction towards the female is of the order of the effective inter-male force.

Following these considerations, we can directly relate the inter-male velocity profile to an effective ’male potential’, *V*_male_ (Fig. 2d). This effective potential describes the averaged effect of an isolated male-male interaction on the dynamics of the males. Under the premise of overdamped motion, the local inter-male velocity between two isolated males is proportional to the local gradient of *V*_male_. To construct *V*_male_, we assume that the effective inter-male forces, which govern the motion of males with respect to each other, depend only on the inter-male distance *d*. This assumption is valid on average, since males rotate to face their opponents when they become engaged with them.

The most basic contribution to *V*_male_ is vibration-based repulsion, which is present throughout the entire range of male-male interaction, and arises from the vibratory threats described above. Since males initially get closer to each other due to their attraction to the female, we expect the vibratory threats to be of shorter range than the attractive logarithmic term of *V*_fem_. Hence we propose that the vibration-based repulsion decays as 1/*d*, much like the dipolar repulsive term of *V*_fem_ (recall Eq. (1)).

The aggression escalates as d decreases beyond a certain average threshold (*d* < 100mm in Fig. 2c). This escalation is mostly channeled through active directed motion of the males towards each other. As the males tend to decrease d when engaged in this type of interaction, it amounts to effective attraction. Fig. 2c suggests that this behaviour should be further dissected into two distinct behavioural regimes, separated by a short-range ’barrier’ (at d ≈ 70mm). We observe that reaching the short-range regime (*d* < 70mm) often entails commitment to grappling. Since these regimes are characterized by relatively short-range linear motion, we model their contributions to *V*_male_ using spring-like terms of the form (*d* − *d*_0_)^2^, where *d*_0_ is an equilibrium distance for the directed motion. These spring-like terms complement the basic vibration-based repulsive term of *V*_male_ when the inter-male distance decreases below *d* ≈ 100mm (Fig. 2d).

We therefore propose to describe *V*_male_ by the piecewise potential

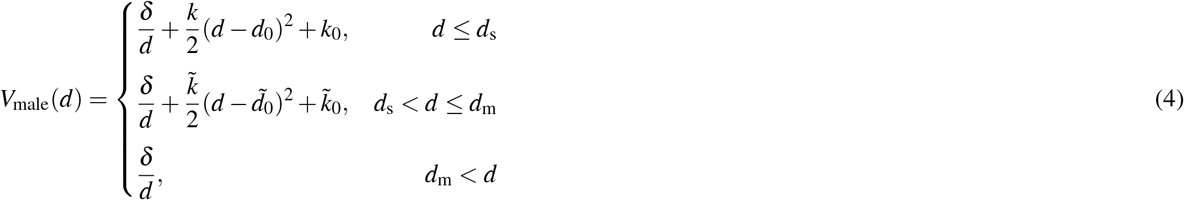

where *δ* > 0 is the strength of the vibration-based repulsion, *k* and 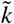 are the effective spring constants for the short-range and medium-range directed motion, respectively, *d*_0_ and 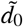 are the corresponding spring equilibrium distances for these two regimes, and *k*_0_, 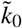 ensure the continuity of the potential at *d* = *d*_s_ ≈ 70*mm* (short-range regime onset) and *d* = *d*_m_ ≈ 100*mm* (medium-range regime onset). Note that the parameters of Eq. (4) are tightly constrained by the shape of the experimental inter-male velocity profile (Fig. 2c). The potential of Eq. (4) and the force profile derived from it are plotted in Fig. 2d,e.

We further note that, according to our observations, when two males take part in a short-range interaction (*d* ≤ *d*_s_) they are completely occupied with each other, and would not resume their approach towards the female until the encounter is resolved. This means that the attraction to the female is effectively ’turned off’ from the perspective of the interacting males when *d* ≤ *d*_s_. To incorporate this behavioural feature in our model, we impose that the attractive term of *V*_fem_ is active only when the males are not engaged in a short-range interaction, so that *V*_fem_ is given only by its repulsive dipole-like term if *d* ≤ *d*_s_, and by Eq. (1) in its entirety if the males are farther apart.

For the full theoretical description of a male in a female’s web, we consider its interactions with the female and with all the other males in the web. The dynamics of male *i* at position **r**_*i*_ = (*r_i_*, *θ_i_*) is governed by its interaction with the effective female potential, given by *V*_fem_(*r_i_*, *θ_i_*, and by its interactions with the effective male potential of any other male *j* at position **r**_*j*_ = (*r_j_*, *θ_j_*), given by *V*_*male j*_(*d_ij_*), where *d_ij_* = |**r**_*i*_ − **r**_*j*_|. Assuming linear superposition of pairwise interactions, male *i* is therefore subjected to a total effective potential, *V*_tot_(*r_i_*, *θ_i_*), that is given by

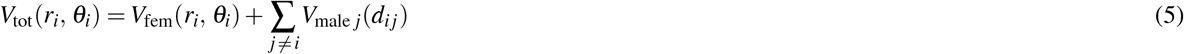

We used Eqs. (1)–(5) to simulate the two-male case (Fig. 2f,g). By adjusting the parameters of Eq. (4), we obtained from these simulations an inter-male velocity profile that is in good agreement with the experimental profile (Fig. 2h). This agreement relies on the already optimized female potential and Langevin parameters, and therefore establishes the effective physics of interacting males in a female’s web. In Supplementary Note 6 we omit the medium-range spring-like regime (*d*_s_ < *d* ≤ *d*_m_) from Eq. (4), and repeat the comparison of Fig. 2h (Supplementary Fig. 6). The results emphasize that this regime is necessary to reproduce the shape of the experimental inter-male velocity profile. In all of the multi-male simulations of this work, the males were initialized on the upper half of a circle that was used to approximate the rim of the orb-web, within which real males can effectively travel continuously (see Fig. 1a; Supplementary Fig. 7). The trajectory sections used to analyze simulation results were only those that remained within this rim-defining circle.

### Interactions between males of different sizes

Thus far, we showed that for interacting males of similar sizes, the dynamics of our model’s ’spider particles’ is in quantitative agreement with the dynamics of real males. However, sexually mature males of the genus *Trichonephila* exhibit an exceptionally high variation of sizes[31, 32]. This poses a long-standing challenge in understanding selection on male body size in these spiders, as males that differ substantially in size are often found to coincide and compete for the same females. To study the role of size in inter-male competition, we have to represent males of different sizes in our model. In the following, we associate the optimized male potential of Fig. 2h with a ’reference’ male, and explore the size-related deviations from it.

The more massive a male is, the stronger the vibrations it can generate and transmit through the web. We account for this effect by varying the strength of the vibration-based repulsive term of *V*_male_, given by the value of δ in Eq. (4). Namely, to represent the interaction of a reference male *i* with a larger male *j*, we set *δ_j_* > *δ*; in their respective male potentials. Now, since *V*_male *j*_(*d_ij_*) > *V*_male *i*_(*d_ij_*), the symmetry of the interaction is broken, as the smaller male feels a more repulsive potential with shallower minima (see Fig. 3a). Note that this property implies that male-male interactions are non-reciprocal.

**Figure 3.**
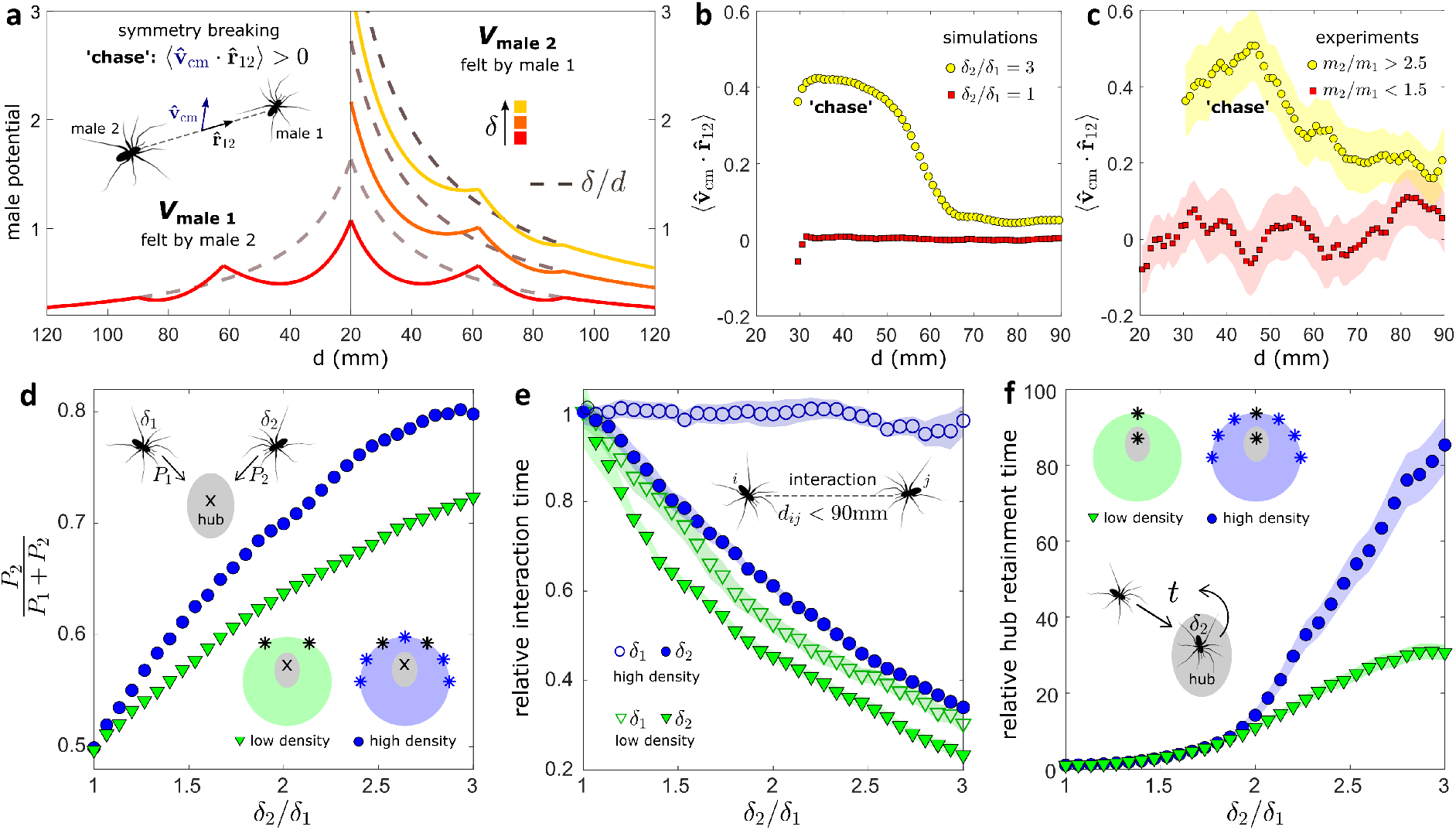
Competitive benefits of large males. **(a)** The strength of the vibration-based repulsive term of *V*_male_, given by *δ*, increases with male size. When a reference male (’male 1’, with *δ*_1_ = *δ*_ref_) interacts with a larger male (’male 2’, with *δ*_2_ > *δ*_1_), the symmetry of the interaction is broken, as the smaller male feels a more repulsive potential with shallower minima. This gives rise to ’chase’ dynamics at small *d*. The ’chase’ is manifested by positive mean correlation between the velocity of the pair’s center of mass 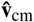, and the inter-male direction vector 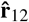, denoted by 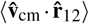. **(b)** Simulated profile of 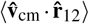. Symmetric case: *δ*_2_/*δ*_1_ = 1, with *δ*_1_ = *δ*_ref_ = 32.7 (*N* = 2000 simulations). Asymmetric case (large size ratio): *δ*_2_/*δ*_1_ = 3, with *δ*_1_ = *δ*_ref_/2 (*N* = 2000 simulations). **(c)** Experimental profile of 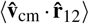. Symmetric case: *m*_2_/*m*_1_ = 1.2(0.1) with *m*_1_ = 12(3)mg and *m*_2_ = 14(4)*mg* (*N* = 19 experiments). Asymmetric case: *m*_2_/*m*_1_ = 2.9(0.3) with *m*_1_ = 4.4(0.9)mg and *m*_2_ = 13(3)*mg* (*N* = 10 experiments). Silhouettes show SEM. In **b** and **c**, a symmetric averaging window of length 11mm was used for smoothing. **(d-f)** Model predictions – effects of male density on large-male advantage. The two probed males, with *δ*_1_ = *δ*_ref_ and *δ*_2_ ≥ *δ*_1_, were initialized as denoted by the black asterisks. The additional males in the high-density setup, with *δ*_i_ = *δ*_ref_, were initialized as denoted by the blue asterisks. **(d)** Relative first arrival probability of the larger ’male 2’. The two probed males have respective first arrival probabilities *P*_1_ and *P*_2_. **(e)** Relative interaction time until arrival to the hub. Two males *i, j* are considered interacting if *d_ij_* < 90mm (within the range of spring-like effective attraction, see **a**). Low and high density setups as in **d**. **(f)** Relative hub retainment time of the larger ’male 2’. In **e** and **f**, each curve is normalized by the respective symmetric case (*δ*_2_/*δ*_1_ = 1).

The symmetry breaking in the interaction of different sized males has a measurable effect on their dynamics, that becomes more prominent if the size difference is large. When a reference male interacts with a larger male, the larger male exerts a stronger effective repulsive force on the reference male, than the reference male does on the larger male. This difference of effective forces, that are exerted in opposite directions along the inter-male axis, gives rise to ’chase’ dynamics at small *d*, where large males tend to chase smaller males away. We can measure the chase in pairs of interacting males by considering the mean correlation between the velocity of the pair’s center of mass, 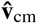, and the inter-male direction vector, 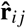, as defined in Fig. 3a. This mean correlation, denoted by 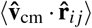, evaluates the extent of asymmetry in the interaction. It is positive and large if significant chase dynamics take place, as predicted for the case of large size difference, and is nearly zero for the symmetric interactions between males of similar sizes (see Fig. 3b).

In our experiments, we find that the average speed of a male on the web increases with its size (Supplementary Fig. 8). To account for this trend in the dynamics of different sized males, the right-hand side of Eq. (2) is scaled by a scaling factor η, given by

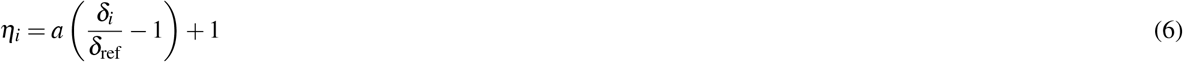

where *η_i_* is the scaling factor for male *i* with vibration-based repulsion strength *δ_i_*, *a* is the fitted slope to the speed-size relation (Supplementary Fig. 8), and *δ*_ref_ is the vibration-based repulsion strength of a reference male (as used in Fig. 2d-h). To obtain Eq. (6), we assumed that *δ_i_* is proportional to the weight of male *i*, so that the ratio between the vibration-based repulsion strengths of two males is simply the ratio of their weights (Supplementary Note 6).

In Fig. 3b,c, we calculated the mean correlation 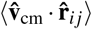 in two-male simulations and experiments, for the case of large size difference, and for the case of similar sizes. The predicted ’chase’ in the case of large size difference, as well as the nearly zero correlation for the symmetric case, clearly appear in both simulations and experiments. This comparison validates our theoretical treatment for males of different sizes, and motivates us to use our model to study realistic scenarios in inter-male competition, in which size may play an important role.

### Large-male advantage and local male density

It is observed that local male density, defined as the number of males occupying a single female’s web, is highly variable in wild populations of *T. clavipes*. A recent study related this observation to the puzzle of high size variability in males, and provided compelling evidence that large males have a significant reproductive advantage when male density is high, and that this advantage is diminished when male density is low[11]. Here we use our model to study the effects of local male density on large-male competitive advantage.

The essential goals of a male in a female’s web is to reach the female’s hub, and retain its hub position until copulation[8, 33]. The competitive advantage of a male in a multi-male setup can therefore be quantified in terms of how often it reaches the hub first (’first arrival’), and how effective it is at retaining its hub position (’hub retainment’). To study how local male density affects large-male advantage in terms of these two measures, we simulated male competition in low-density and high-density setups, as illustrated in the insets of Fig. 3d,f.

First, we explore the outcomes of competition with respect to first arrival (Fig. 3d,e). We simulated the competition between two probed males that enter the female’s web at the same time and at spatially equivalent positions (marked by black asterisks in the bottom insets of Fig. 3d). In the low-density setup, the system only includes these two competing males. In the high-density setup, we added five more males of the reference size. The additional males were initialized in a symmetric configuration (marked by blue asterisks), such that the initial positions of the probed males remained equivalent. We note that the symmetry of the initial configurations quickly disappears due to stochastic effects, and is only used to ensure that there is no bias in the simulations. Of the two probed males, one is of the reference size (’male 1’, with *δ*_1_ = *δ*_ref_), and the other of a variable size (’male 2’, with *δ*_2_ ≥ *δ*_1_). We increased the size of ’male 2’ in small increments, starting from the reference size, and up to 3-times the reference size.

In Fig. 3d, we plot the relative probability of the larger ’male 2’ (relative to the reference ’male 1’) to arrive first at the female’s hub, as a function of the size ratio between the two males. We find that the first arrival probability of the larger male increases with its size. This follows from the faster speed of the larger male, and its ability to chase the smaller competitor away, if they happen to encounter each other. However, even for a large size ratio of 3, the smaller probed male still has a first arrival probability of ~ 30% in the low-density setup. In the high-density setup, we find that the relative first arrival probability of the larger male is enhanced compared to the low density case, in agreement with the empirical findings[11].

When male density is high, males frequently encounter rivals and interact with them. We demonstrate how the abundance of male-male interactions creates a competitive environment that is particularly advantageous for large males. For this analysis, two males i, *j* are considered interacting if *d_ij_* is within the range of spring-like effective attraction (*d_ij_* < 90mm, see Fig. 3a). In Fig. 3e, we plot the mean time spent in male-male interactions by each of the probed males of Fig. 3d, during their journey from the edge of the female’s web, and until they reach the hub. In the low-density setup, we find that this interaction time is similar for both males, and that it decreases with the size of the larger male. This shows that the stronger vibration-based repulsion of large males helps them avoid long-lasting short-range interactions, as they typically chase smaller males away. The ability to avoid short-range interactions becomes significant in the high-density setup. We find that increased interactions with the additional males delay the reference-sized ’male 1’ substantially, while the larger ’male 2’ remains effective at avoiding these interactions (see also Supplementary Fig. 9, which visualizes this phenomenon in spatial interaction maps).

Finally, we study the competitive advantage of large males with respect to hub retainment. We initialized the simulations with the larger ’male 2’ already in the female’s hub (the global minimum of *V*_fem_), and the remaining reference-sized males entering the web in a symmetric configuration (see Fig. 3f insets). In Fig. 3f, we plot the mean time until ’male 2’ has lost its hub position, and been replaced by any of the other males, as a function of the size ratio. This time is normalized by its value in the symmetric case (*δ*_1_ = *δ*_2_ = *δ*_ref_ in the respective setup). We find that the hub retainment of the larger male increases dramatically with its size. For a size ratio of 3 in the low-density setup, the hub retainment time of the larger male is ~ 30 times longer than that of a reference-sized male. In the high-density setup, this relative advantage is greatly pronounced. This shows that large males remain effective at retaining their hub position in high male densities, while the abundance of male-male interactions penalize small males even if they managed to obtain the hub position. Note that even large males have a finite hub retainment time, which saturates in the low-density case, due to the random component of the agent’s motion.

## Discussion

Here we formulated a quantitative model of spider behaviour during mate search and competition, based on effective physical forces. This approach considers spiders as particles responding only to effective forces, and assumes no memory, cognition, or decision-making on the part of the agents. Despite the simplicity of the agents, there is strong quantitative agreement between the dynamical characteristics of the particles in the model and the movement and interactions of real male spiders. Our model and associated empirical observations are consistent with Morgan’s Canon, stating that “in no case is an animal activity to be interpreted in terms of higher psychological processes if it can be fairly interpreted in terms of processes which stand lower in the scale of psychological evolution and development”[34]. Our results demonstrate that a minimal behavioural repertoire of responding to seismic and geometric information is all that is required from the male spiders in order to solve the tasks they are faced with on the web. This is an important extension of insight into the processes underlying emergent properties of organismal interactions. While fields such as collective animal behaviour have a long history of revealing simple rules that can recreate complex collective level phenomena[35, 36], to our knowledge this is the first such example in the context of mating behaviour and male-male competition.

The first task of the reproductively foraging male spider is to locate a female’s web. In *Trichonephila*, pheromonal cues are used by males to both locate and select among females, with spiders commonly taking these cues directly from the silk itself[37]. However, once males have entered the orb web, they rely primarily on seismic information to interact with the female, as well as with rival males[19, 22]. We find that upon entering the orb, males move around the perimeter of the web, thereby maintaining the greatest possible distance to the dangerous female, until they are in a position above and behind the hub where the female resides.

While on the female’s web, males are likely to encounter competition from rival males, either from those who are already present, or those that arrive later[12]. Because males of this genus form waiting queues in which proximity to the female influences the order in which they will mate[38], and therefore their chances of reproductive success, it can be advantageous to fight and displace rival males. Although males themselves are unlikely to injure one another in these fights, conflict between males creates vibrational disturbances on the web that can lead to attack by the larger female, so male-male competition is inherently risky[39]. Because of this risk, physical contact between males is preceded by a period of signalling and assessment, which we modelled in the form of an effective double-well potential. Here again we showed that a seemingly complex interaction, which in many other taxa is assumed to involve higher order cognitive and reasoning processes, can be successfully accounted for by a modelling approach that relies solely on effective physical interactions.

Moreover, in spiders and many other taxa, the outcome of male-male contests is strongly associated with size[40], and here we showed that these dynamics can also be recaptured by purely physical models that assume no memory, reasoning, or other cognitive assessment of the rival’s ability. Furthermore, based solely on pairwise male-male interactions, our model is able to capture competitive dynamics in webs containing many males. Namely, based on the same simple interaction rules, the model explains the previously observed increase of large-male advantage with male density[11], as well as advantages associated with prior web residence in this genus[12]. We have shown in our simulations that the mechanism responsible for these phenomena is related to the ability of large males to avoid long-lasting male-male interactions, and this prediction remains to be explored in future experiments.

The ability to locate and compete for mates is a behaviour with clear fitness consequences. Our results suggest that in male spiders, selection has generated an efficient solution to this complex task, potentially relying almost entirely on seismic information. This is in some ways not surprising – once on the female web, male spiders exist in a seismic world, and so should rely heavily on this modality[20]. Furthermore, the small size and constrained behavioural repertoire (no prey capture, no web-building) of males in this genus may limit the capacity for advanced cognition and reasoning ([17], but see [41] for an argument for extended cognition in spiders using the web). Combined with the incredibly strong selective pressure exerted by male-male competition, and the risks posed by the larger female, there is ample opportunity for natural selection to produce an elegant solution to these problems. Nevertheless, it is striking that the ostensibly complex problem of mate search and male competition in this species may be solved by biological agents employing rules no more sophisticated than the interactions among physical particles.

## Methods

### Experimental setup and analysis of trajectories

Experiments were carried out in an experimental area in Gamboa, Panama, with female and male *Trichonephila clavipes*. We collected spiders from Barro Colorado Island (BCI), Panama (9.15°N, 79.85°W) in April 2018, April 2019 and September 2019, and transferred them to an experimental area in Gamboa, Panama (9.12°N, 79.70°W). We placed females on 70 × 70cm timber or metal frames, where they could build their webs (Supplementary Fig. 1a), and kept males in individual plastic containers. Females built their webs overnight, and experiments were conducted the following day with females that had built webs entirely within the frames (as in Supplementary Fig. 1a). Webs were sprayed with water once a day, and a single cricket fed to female spiders three times a day. One of these feeding sessions occurred immediately before recording began to reduce the risk of death by an aggressive female for experimental males.

For the behavioural experiments with a single male, we placed males at initial positions near the rim of the orb-web, such that the entire set of initial positions covers all of the peripheral areas of the web – in order to gauge the dynamics of males on the entire surface of the web (Supplementary Fig. 2). In experiments with two males, we placed the males simultaneously in the upper half of the web (above the down-facing female), and roughly at spatially equivalent positions on either sides of the female (as in Supplementary Fig. 1b). We recorded the male and female movement on the web against a white background sheet using a Sony A7s II (Supplementary Fig. 1b). All experimental work was conducted in compliance with the Smithsonian Tropical Research Institute.

Using the object detection algorithm Mask-RCNN, we obtained the trajectories from the behavioural recordings by training the network on a subset of manually annotated video screenshots to accurately detect individual spiders (for details see [42]). This resulted in a pixel mask for each detected spider, where the mask’s centroid was computed as the individuals’ position at that frame. These positions were then used to automatically reconstruct continuous spider trajectories using a distance-based identity assignment approach. Lastly, we used TrackUtil for manual track corrections, such as keeping a unique identity for each individual. To standardize the raw tracks, we referenced them to the same coordinate frame through shifting and scaling (from pixel to cm), resulting in the four corners of the frame to match.

## Supporting information

caption for Supplementary Movie 1

## References

[1] Michael J Ryan and A Stanley Rand. “Female responses to ancestral advertisement calls in túngara frogs”. In: Science 269.5222 (1995), pp. 390–392.

[2] A Stanley Rand, Michael J Ryan, and Walter Wilczynski. “Signal redundancy and receiver permissiveness in acoustic mate recognition by the túngara frog, Physalaemus pustulosus”. In: American Zoologist 32.1 (1992), pp. 81–90.

[3] Aviram Gelblum et al. “Ant groups optimally amplify the effect of transiently informed individuals”. In: Nature communications 6 (2015), p. 7729.

[4] Ofer Feinerman et al. “The physics of cooperative transport in groups of ants”. In: Nature Physics 14.7 (2018), p. 683.

[5] Madeleine Beekman and Tanya Latty. “Brainless but multi-headed: decision making by the acellular slime mould Physarum polycephalum”. In: Journal of molecular biology 427.23 (2015), pp. 3734–3743.

[6] Chris R Reid and Tanya Latty. “Collective behaviour and swarm intelligence in slime moulds”. In: FEMS microbiology reviews 40.6 (2016), pp. 798–806.

[7] Leann Myers and Terry Christenson. “Transition from predatory juvenile male to mate-searching adult in the orb-weaving spider Nephila clavipes (Araneae, Araneidae)”. In: The Journal of Arachnology 16.2 (1988), pp. 254–257.

[8] Terry E Christenson and Kenneth C Goist. “Costs and benefits of male-male competition in the orb weaving spider, Nephila clavipes”. In: Behavioral Ecology and Sociobiology 5.1 (1979), pp. 87–92.

[9] JM Schneider and MA Elgar. “Sexual cannibalism in Nephila plumipes as a consequence of female life history strategies”. In: Journal of Evolutionary Biology 15.1 (2002), pp. 84–91.

[10] Matthias W Foellmer and Daphne J Fairbairn. “Males under attack: sexual cannibalism and its consequences for male morphology and behaviour in an orb-weaving spider”. In: Evolutionary Ecology Research 6.2 (2004), pp. 163–181.

[11] Clare C Rittschof. “Male density affects large-male advantage in the golden silk spider, Nephila clavipes”. In: Behavioral Ecology 21.5 (2010), pp. 979–985.

[12] Lyndon Alexander Jordan, Hanna Kokko, and Michael Kasumovic. “Reproductive foragers: male spiders choose mates by selecting among competitive environments”. In: The American Naturalist 183.5 (2014), pp. 638–649.

[13] Graham H Pyke, H Ronald Pulliam, and Eric L Charnov. “Optimal foraging: a selective review of theory and tests”. In: The quarterly review of biology 52.2 (1977), pp. 137–154.

[14] Graham H Pyke. “Optimal foraging theory: a critical review”. In: Annual review of ecology and systematics 15.1 (1984), pp. 523–575.

[15] Joel S Brown, John W Laundré, and Mahesh Gurung. “The ecology of fear: optimal foraging, game theory, and trophic interactions”. In: Journal of mammalogy 80.2 (1999), pp. 385–399.

[16] Lars Chittka and Jeremy Niven. “Are bigger brains better?” In: Current biology 19.21 (2009), R995–R1008.

[17] Jeremy E Niven and Sarah M Farris. “Miniaturization of nervous systems and neurons”. In: Current Biology 22.9 (2012), R323–R329.

[18] Robert R Jackson and Fiona R Cross. “Spider cognition”. In: Advances in insect physiology. Vol. 41. Elsevier, 2011, pp. 115–174.

[19] Marie Elisabeth Herberstein. Spider behaviour: flexibility and versatility. Cambridge University Press, 2011.

[20] MA Landolfa and FG Barth. “Vibrations in the orb web of the spider Nephila clavipes: cues for discrimination and orientation”. In: Journal of Comparative Physiology A 179.4 (1996), pp. 493–508.

[21] Todd A Blackledge, Matjaž Kuntner, and Ingi Agnarsson. “The form and function of spider orb webs: evolution from silk to ecosystems”. In: Advances in insect physiology. Vol. 41. Elsevier, 2011, pp. 175–262.

[22] Marie E Herberstein et al. “Dangerous mating systems: signal complexity, signal content and neural capacity in spiders”. In: Neuroscience & Biobehavioral Reviews 46 (2014), pp. 509–518.

[23] Samuel Zschokke and Kensuke Nakata. “Spider orientation and hub position in orb webs”. In: Naturwissenschaften 97.1 (2010), p. 43.

[24] Thomas Krakauer. “Thermal responses of the orb-weaving spider, Nephila clavipes (Araneae: Argiopidae)”. In: American Midland Naturalist (1972), pp. 245–250.

[25] Terry E Christenson et al. “Mating behavior of the golden-orb-weaving spider, Nephila clavipes: I. Female receptivity and male courtship.” In: Journal of comparative psychology 99.2 (1985), p. 160.

[26] Samuel Zschokke et al. “Form and function of the orb-web”. In: European arachnology 19 (2000), p. 99.

[27] CG Joslin and CG Gray. “Multipole expansions in two dimensions”. In: Molecular Physics 50.2 (1983), pp. 329–345.

[28] Pawel Romanczuk et al. “Active brownian particles”. In: The European Physical Journal Special Topics 202.1 (2012), pp. 1–162.

[29] Giorgio Volpe, Sylvain Gigan, and Giovanni Volpe. “Simulation of the active Brownian motion of a microswimmer”. In: American Journal of Physics 82.7 (2014), pp. 659–664.

[30] Jeffrey Cohn, Frances V Balding, and Terry E Christenson. “In defense of Nephila clavipes: Postmate guarding by the male golden orb-weaving spider.” In: Journal of Comparative Psychology 102.4 (1988), p. 319.

[31] Fritz Vollrath. “Male body size and fitness in the web-building spider Nephila clavipes”. In: Zeitsch-rift fir Tierpsychologie 53.1 (1980), pp. 61–78.

[32] JM Schneider et al. “Sperm competition and small size advantage for males of the golden orb-web spider Nephila edulis”. In: Journal of Evolutionary Biology 13.6 (2000), pp. 939–946.

[33] Clovis W Moore. “The life cycle, habitat and variation in selected web parameters in the spider, Nephila clavipes Koch (Araneidae)”. In: American Midland Naturalist (1977), pp. 95–108.

[34] Elliott Sober. “Morgan’s canon.” In: (1998).

[35] Tamás Vicsek et al. “Novel type of phase transition in a system of self-driven particles”. In: Physical review letters 75.6 (1995), p. 1226.

[36] Iain D Couzin et al. “Collective memory and spatial sorting in animal groups”. In: Journal of theoretical biology 218.1 (2002), pp. 1–11.

[37] Gabriele Uhl. “Spider olfaction: attracting, detecting, luring and avoiding”. In: Spider ecophysiology. Springer, 2013, pp. 141–157.

[38] Mark A Elgar. “Sexual cannibalism, size dimorphism, and courtship behavior in orb-weaving spiders (Araneidae)”. In: Evolution 45.2 (1991), pp. 444–448.

[39] Jutta M Schneider and Mark A Elgar. “Sexual cannibalism and sperm competition in the golden orb-web spider Nephila plumipes (Araneoidea): female and male perspectives”. In: Behavioral Ecology 12.5 (2001), pp. 547–552.

[40] Gareth Arnott and Robert W Elwood. “Assessment of fighting ability in animal contests”. In: Animal Behaviour 77.5 (2009), pp. 991–1004.

[41] Hilton F Japyassú and Kevin N Laland. “Extended spider cognition”. In: Animal Cognition 20.3 (2017), pp. 375–395.

[42] Armando Levid Rodriguez-Santiago et al. “A Simple Methodology for 2D Reconstruction Using a CNN Model”. In: Mexican Conference on Pattern Recognition. Springer. 2020, pp. 98–107.

