## Supplementary Information for "Dynamics on the web: spiders use physical rules to solve complex tasks in mate search and competition"

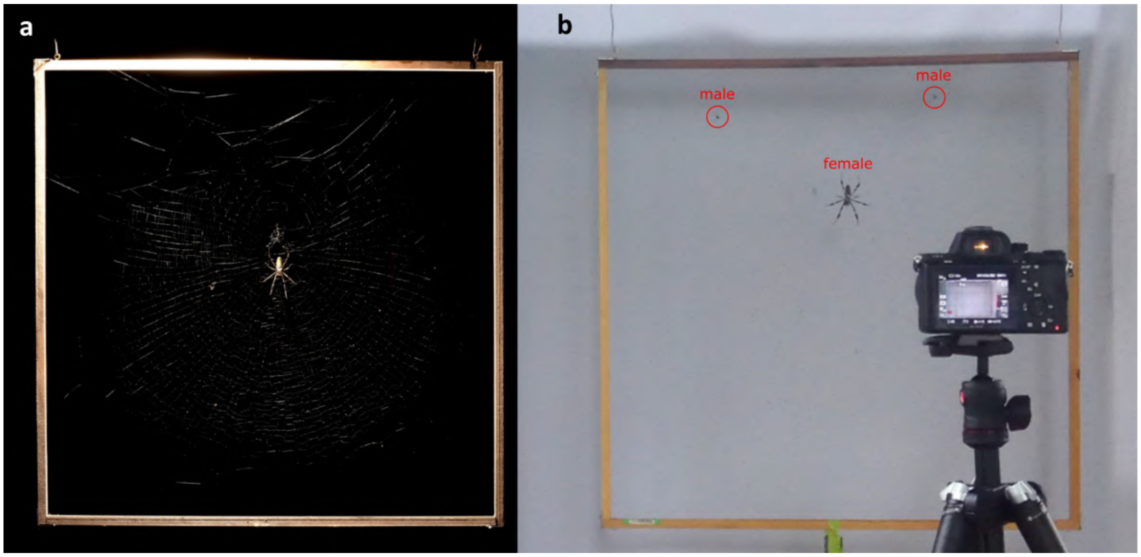


**Supplementary Figure 1.** Experimental setup. **(a)** Individual females were placed on a $70 \times 70cm$ timber or metal frames, where they would build their webs overnight. Only females that had built their web entirely within the frame (as in the picture) were used for the experiments. **(b)** The movement of the spiders during each experiment was recorded against a white background sheet using a Sony A7s II. The picture is a snapshot from the beginning of a two-male experiment. In these experiments, the males were initially placed simultaneously in the upper half of the web, and roughly at spatially equivalent positions on either sides of the female, as in the picture.

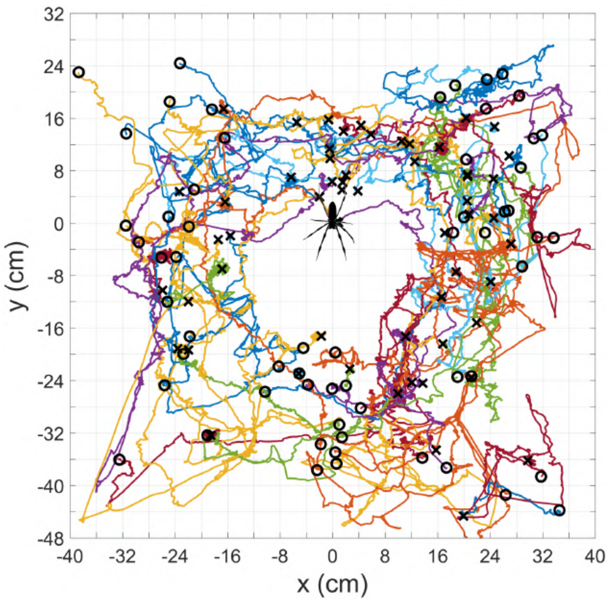


**Supplementary Figure 2.** Experimental single-male trajectories ($N = 59$). The individual trajectories are superimposed according to the female-centered coordinate system, such that the position of the female's head is the point of reference. In each of these experiments, we placed a single male at some initial position near the rim of the orb-web, such that the entire set of initial positions covers all of the peripheral areas of the web, as depicted by the black circles. The left-right symmetry (a lack of left-right bias) in this set of trajectories can be observed. Each trajectory was obtained from about 10 minutes of single-male dynamics. Black x's denote final positions.

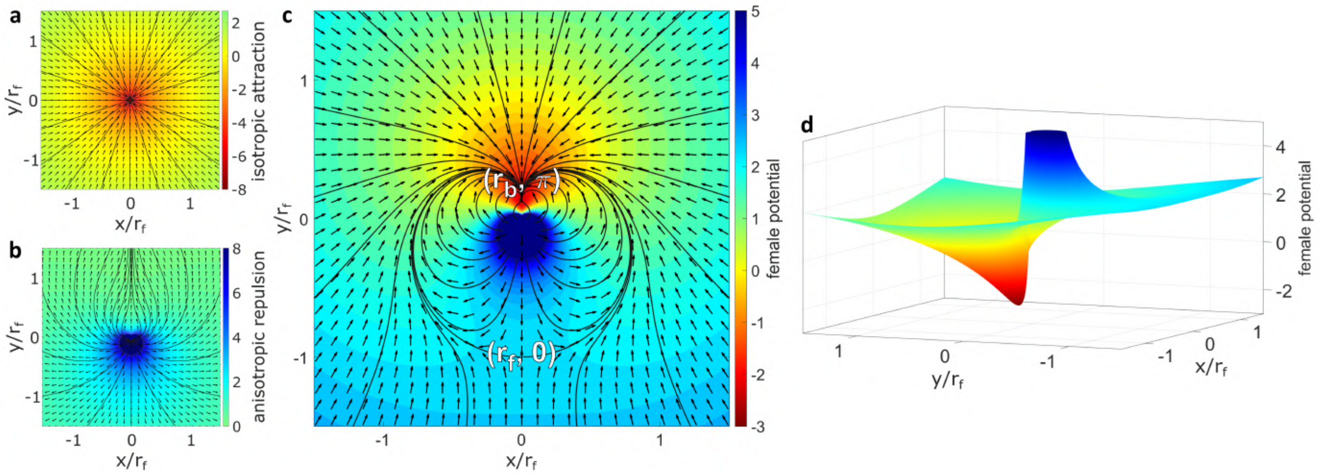


**Supplementary Figure 3.** Visualisation of the female effective potential. **(a, b)** The attractive (isotropic) and repulsive (anisotropic) components of the female effective potential, respectively. **(c)** The overall female effective potential, given by the sum of **a** and **b** (Eq. (S1)). This potential has, in polar coordinates, a global minimum at $(r_\mathrm{b}, \pi)$, and an unstable saddle at $(r_\mathrm{f}, 0)$. In **a**-**c**, the contours (indicated by the color-map) of the potentials are overlaid with their corresponding flow fields (arrows) and stream lines (black lines). **(d)** A 3-dimensional representation of the potential landscape of **c**. In **a** and **b**, the divergence at the origin is truncated.

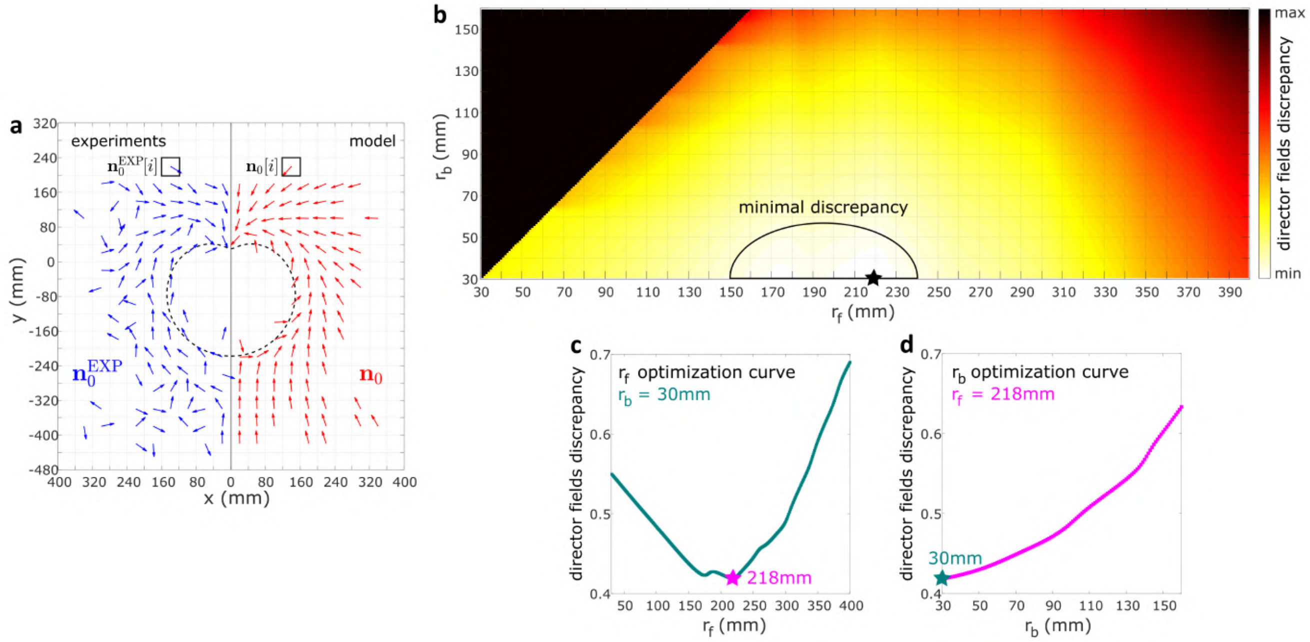


**Supplementary Figure 4.** Optimization of the female effective potential. **(a)** The director field of the female potential ($\mathbf{n}_0$) is shown alongside its experimental analogue ($\mathbf{n}_0^{\mathrm{EXP}}$). Two 'corresponding' bins, $\mathbf{n}_0^{\mathrm{EXP}}[i]$ and $\mathbf{n}_0[i]$ (as denoted in Eq. (S14)), are shown for clarity. **(b)** A heat map visualisation of the female potential's parameters optimization. The color bar indicates the value of the mean discrepancy between $\mathbf{n}_0$ and $\mathbf{n}_0^{\mathrm{EXP}}$ for different choices of $(r_\mathrm{b}, r_\mathrm{f})$. A visual estimate for the subset of $(r_\mathrm{b}, r_\mathrm{f})$ values which yielded the lowest discrepancy, as well as the actual minimum, are marked by a half-ellipse and a star, respectively. **(c, d)** One-dimensional optimization curves for $r_\mathrm{f}$ at the optimal $r_\mathrm{b}$, and for $r_\mathrm{b}$ at the optimal $r_\mathrm{f}$, respectively. The minimum of each curve is marked by a star.

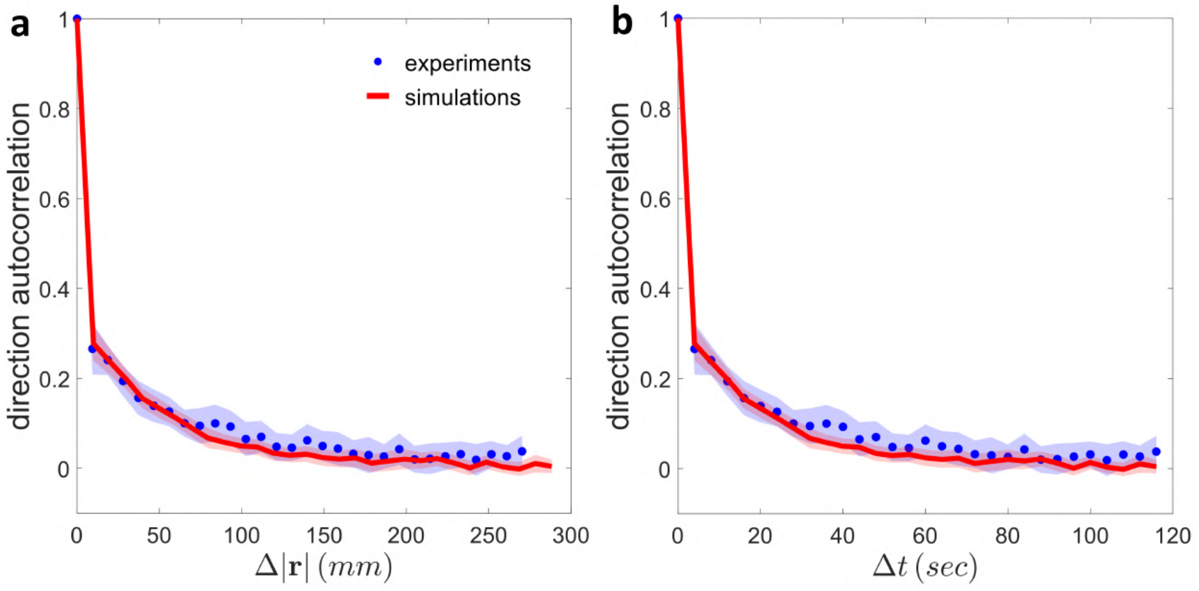


**Supplementary Figure 5.** Autocorrelation of the direction of movement along male trajectories with respect to **(a)** distance ($\Delta|\mathbf{r}|$), and **(b)** time ($\Delta t$). The experimental trajectories were obtained from the single-male experiments shown in Supplementary Fig. 2. The simulated trajectories were obtained from single-male simulations according to Eqs. (S15) and (S16), with the optimized female potential ($V = \bar{V}_{\text{fem}}$ with $r_\text{b} = 30\,mm$, $r_\text{f} = 218\,mm$, and $C = 1.32$), and Langevin parameters $\gamma = 2/sec$, $v_\text{p} = 2\,mm/sec$, $D_T = 5\,mm^2/sec$, and $D_R = 0.05\,rad^2/sec$. The simulated single-male trajectories were produced for the same initial positions and durations as the experimental ones. Silhouettes show SEM.

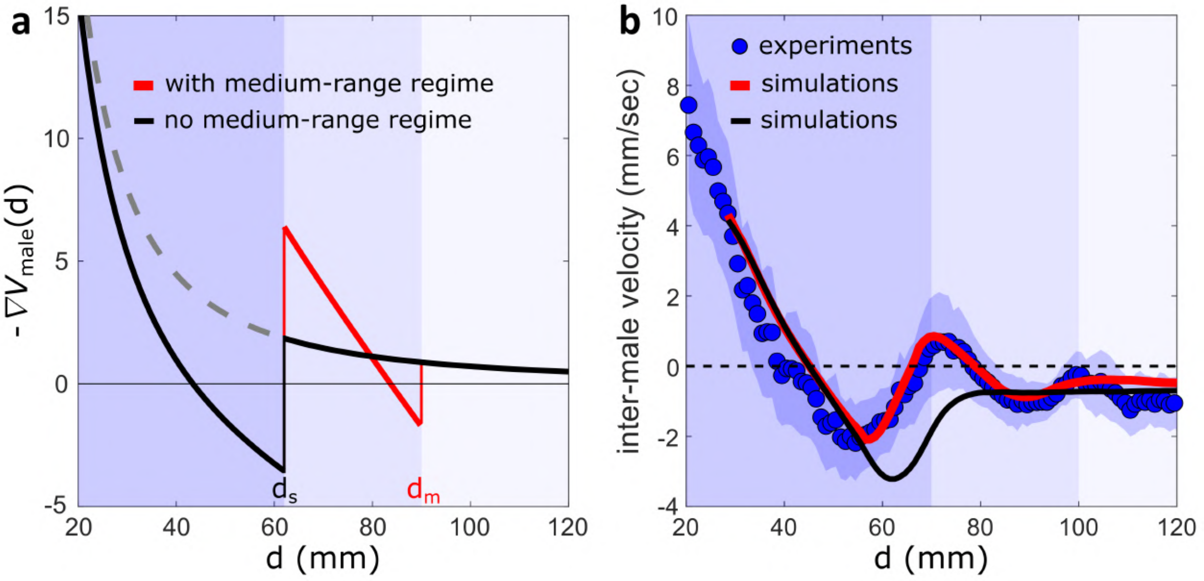


**Supplementary Figure 6.** **(a)** Comparison of the force profiles derived from $V_{\text{male}}$ with and without the medium-range spring-like regime. These force profiles differ only in the range $d_\text{s} < d \leq d_\text{m}$. The grey dashed line extrapolates the vibration-based repulsive term ($\delta/d^2$). **(b)** Inter-male velocity profiles comparison. The red and black profiles were obtained from simulations ($N = 2000$) with the male potential as in Eq. (S18) and Eq. (S17), respectively. A symmetric averaging window of length $11mm$ was used for smoothing.

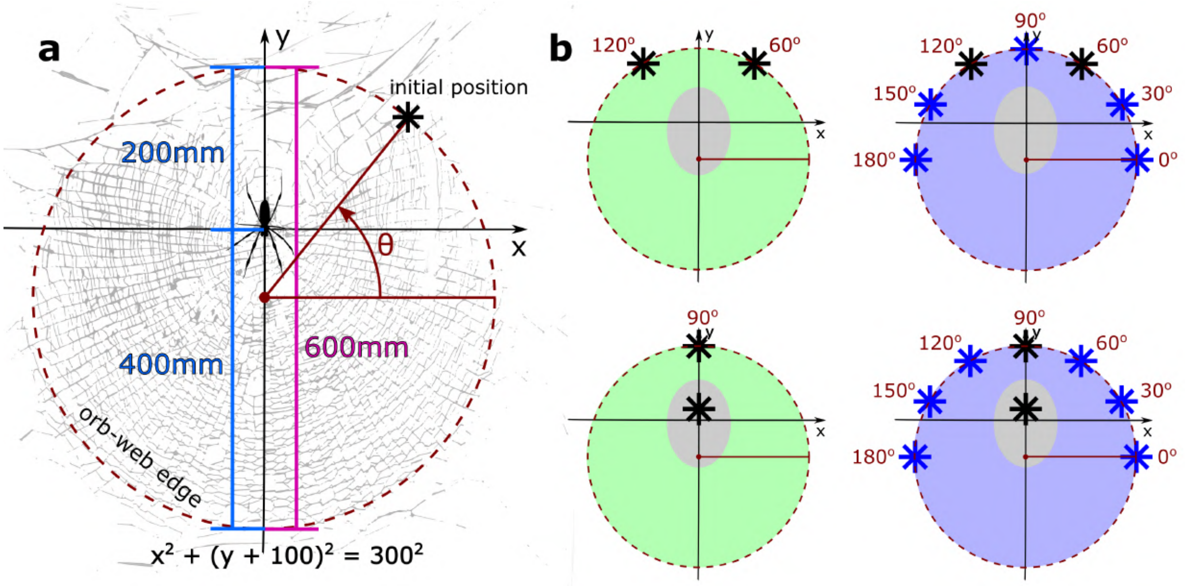


**Supplementary Figure 7.** **(a)** The circle used to approximate the rim of the orb-web, within which real males can effectively travel continuously. In accordance with the geometry of a real web, the center of the rim-defining circle is located below the head of the down-facing female, that is below the origin of the female-centered coordinate system. The dimensions of the circle and its position, as defined by the circle's equation, were chosen to approximate the rim of an adult female's web, but note that the dimensions of these webs can vary substantially. Note that in our simulations, the movement of males is **not** restricted to the inside of this circle. In all of the multi-male simulations of this work, the males were initialized on the upper half of the circle. Note that the angle $\theta$ is defined here with respect to the circle's center, and is **not** the same angle that describes the position of a male in the female-centered coordinate system. Here, we use this angle to easily describe the initial positions in the multi-male configurations. **(b)** Initial positions of males in terms of the angle $\theta$, as defined in **a**, in the multi-male configurations used to simulate low and high male density.

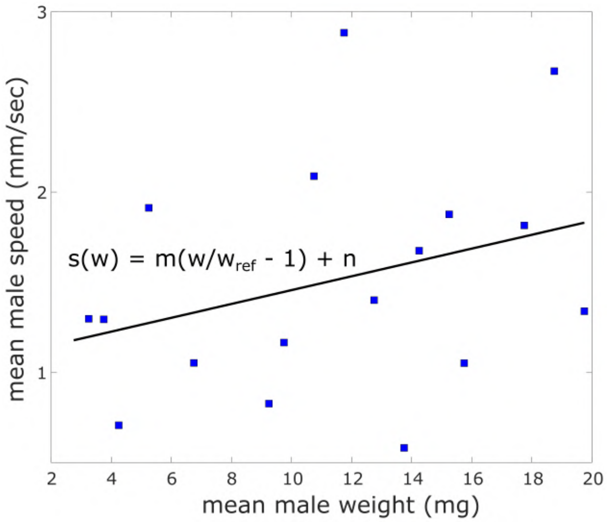


**Supplementary Figure 8.** The average speed of males on the web as a function of their size (in terms of their weight). The line is the fit to the shown linear equation.

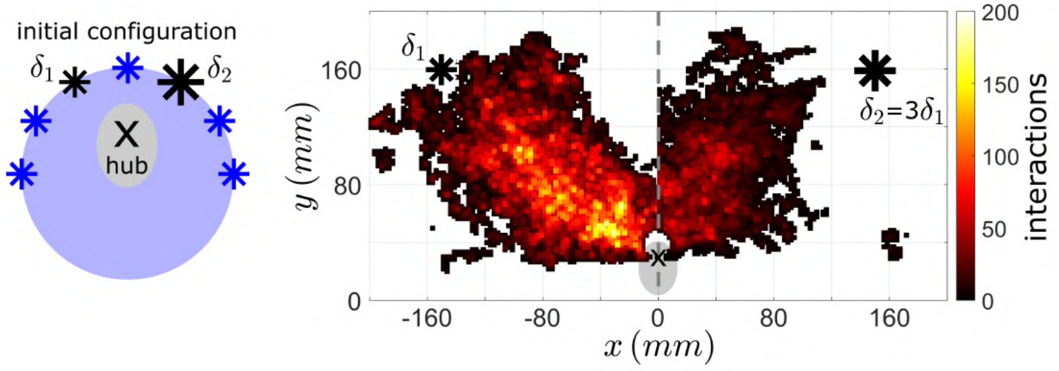


**Supplementary Figure 9.** Spatial interactions map of each of the probed males in the high-density setup, for the largest simulated size ratio (the vibration-based repulsion strength of 'male 2' is 3-times that of 'male 1', that is $\delta_2 = 3\delta_1$). For this analysis, two males $i$, $j$ are considered interacting if $d_{ij} < 90mm$. Each data point corresponds to a $1 \times 1mm$ position bin. The color of each data point indicates the number of independent simulations in which a probed male participated in an interaction while passing through its bin. Out of 200 first arrival cases for each male, we see that the map of the larger male features significantly less interactions. For each probed male, only the interactions that occurred in the half-plane in which it was initialized are shown, to avoid clutter. Asterisks mark initial positions.

### Supplementary Note 1: Fixed points of the female potential's flow-field

Recall the female effective potential given in Eq. (1) of the main text,

$$V_{\text{fem}}(r,\theta) = \alpha \ln r + \beta \frac{C + \cos\theta}{r} \tag{S1}$$

The effective force derived from Eq. (S1), given by $-\nabla V_{\text{fem}}$, generates a flow-field with two fixed points. Here we obtain these fixed points and their stability. The radial and angular partial derivatives of Eq. (S1) are

$$\frac{\partial V_{\text{fem}}}{\partial r} = \alpha \frac{1}{r} - \beta \frac{C + \cos\theta}{r^2} \tag{S2}$$

$$\frac{\partial V_{\text{fem}}}{\partial \theta} = -\beta \frac{\sin\theta}{r} \tag{S3}$$

Eq. (S3) is equal to zero when $\theta = 0$ (in front of the female) and when $\theta = \pi$ (behind the female). Plugging each of these values of $\theta$ into Eq. (S2), setting it to zero, and solving for $r$, we find the two fixed points of Eq. (S1),

$$(r_{\text{f}}, 0) = \left(\frac{\beta}{\alpha}(C+1), 0\right), \qquad (r_{\text{b}}, \pi) = \left(\frac{\beta}{\alpha}(C-1), \pi\right). \tag{S4}$$

The locations of these fixed points are shown in Supplementary Fig. 3c. The flow field shows that $(r_{\text{f}}, 0)$ is an unstable saddle point, and that $(r_{\text{b}}, \pi)$ is a global minimum. The unstable saddle point is associated with the boundary of the frontal sector directly in front of the down-facing female, and the globally stable fixed point is associated with the location of the hub (Fig. 1a in main text).

### Supplementary Note 2: Rescaling the female potential

We wish to rescale $V_{\text{fem}}$ and express its parameters in terms of the natural scales $r_{\text{b}}$ and $r_{\text{f}}$. Since these lengths have experimental analogous that can be estimated from the average flow-field of males around females, the rescaling will allow us to estimate the parameters of $V_{\text{fem}}$. By dividing the expression for $r_{\text{b}}$ by the expression for $r_{\text{f}}$ (Eq. (S4)) and solving for $C$, we get

$$C = \frac{1 + r_{\text{b}}/r_{\text{f}}}{1 - r_{\text{b}}/r_{\text{f}}} \tag{S5}$$

We see that the value of the constant $C$ depends explicitly only on the ratio $r_{\text{b}}/r_{\text{f}}$. Rearranging the expression for $r_{\text{f}}$ in Eq. (S4), we get a relation that will be used to rescale $V_{\text{fem}}$,

$$r_{\text{f}} \frac{\alpha}{\beta} = C + 1 \tag{S6}$$

We now rescale the potential in Eq. (S1) by defining

$$\bar{V}_{\text{fem}} = \frac{r_{\text{f}}}{\beta} V_{\text{fem}} - (C+1)\ln(r_{\text{f}}) \tag{S7}$$

Using Eqs. (S1), (S6) and (S7), we get the rescaled dimensionless potential $\bar{V}_{\text{fem}}$,

$$\bar{V}_{\text{fem}} = \frac{r_{\text{f}}}{\beta}\left(\alpha \ln r + \beta \frac{C+\cos\theta}{r}\right) - (C+1)\ln(r_{\text{f}}) = (C+1)\ln(r/r_{\text{f}}) + \frac{C+\cos\theta}{(r/r_{\text{f}})} \qquad \Rightarrow$$

$$\bar{V}_{\text{fem}}(\bar{r},\theta) = (C+1)\ln(\bar{r}) + \frac{C+\cos\theta}{\bar{r}} \tag{S8}$$

where $\bar{V}_{\text{fem}}$ is now a function of the rescaled dimensionless distance $\bar{r} = r/r_{\text{f}}$.

### Supplementary Note 3: Female potential's parameters estimation

To estimate the parameters of $V_{\text{fem}}$, we compare the normalized flow-field (director field) of $V_{\text{fem}}$, given by $-\nabla V_{\text{fem}}/|\nabla V_{\text{fem}}|$, with its experimental analogue, obtained from single-male trajectories (Supplementary Fig. 4a). The gradient of the rescaled female potential $\bar{V}_{\text{fem}}$ is given by

$$\nabla \bar{V}_{\text{fem}}(\bar{r},\theta) = \frac{\partial \bar{V}_{\text{fem}}}{\partial \bar{r}}\hat{r} + \frac{1}{\bar{r}}\frac{\partial \bar{V}_{\text{fem}}}{\partial \theta}\hat{\theta} = -(\bar{F}_r \hat{r} + \bar{F}_\theta \hat{\theta}) \tag{S9}$$

where $\bar{F}_r$ and $\bar{F}_\theta$ are the magnitudes of the forces due to $\bar{V}_{\text{fem}}$ along the $\hat{r}$ and $\hat{\theta}$ directions, respectively. They are given by

$$\bar{F}_r(\bar{r},\theta) = \frac{C+1}{\bar{r}} - \frac{C+\cos\theta}{\bar{r}^2}, \qquad \bar{F}_\theta(\bar{r},\theta) = -\frac{\sin\theta}{\bar{r}^2} \tag{S10}$$

In Cartesian coordinates, the forces due to $\bar{V}_{\text{fem}}$ are related to $\bar{F}_r$ and $\bar{F}_\theta$ through

$$\bar{F}_x = \bar{F}_r\cos\theta - \bar{F}_\theta \sin\theta, \qquad \bar{F}_y = \bar{F}_r \sin\theta + \bar{F}_\theta\cos\theta \tag{S11}$$

For overdamped motion, the velocities are directly proportional to the forces, so we can write

$$\mathbf{v}_0 = \gamma^{-1}\begin{pmatrix}\bar{F}_x\\ \bar{F}_y\end{pmatrix} \tag{S12}$$

where $\mathbf{v}_0 = \mathbf{v}_0(\bar{r},\theta)$ is the deterministic component of a male's velocity field due only to $\bar{V}_{\text{fem}}$, and $\gamma$ is the effective friction coefficient associated with the motion of males on a female's web. For comparison of the deterministic flow due to $\bar{V}_{\text{fem}}$ with its experimental analog, we consider only the direction of $\mathbf{v}_0$,

$$\mathbf{n}_0 = \frac{\mathbf{v}_0}{|\mathbf{v}_0|} \tag{S13}$$

where $\mathbf{n}_0$ is the director field (normalized velocity field) due to $\bar{V}_{\text{fem}}$.

We now describe the optimization procedure used to obtain estimates for $r_{\text{b}}$ and $r_{\text{f}}$. We obtain an experimental director field, $\mathbf{n}_0^{\text{EXP}}$, from the data set of single-male trajectories, by calculating the mean direction of male movement in small bins (see Supplementary Fig. 4a). With the same resolution (same bin size), we obtain the theoretical director field of $\bar{V}_{\text{fem}}$ (Eq. (S13)). The mean discrepancy (MD) between $\mathbf{n}_0^{\text{EXP}}$ and $\mathbf{n}_0$ is calculated for different choices of $(r_{\text{b}}, r_{\text{f}})$ as

$$\text{MD}(r_{\text{b}}, r_{\text{f}}) = \frac{\sum_{i=1}^{N_{\text{bin}}} |\mathbf{n}_0^{\text{EXP}}[i] - \mathbf{n}_0[i](r_{\text{b}}, r_{\text{f}})|}{N_{\text{bin}}} \tag{S14}$$

where $i$ is the (arbitrary) index of spatially equivalent bins, $\mathbf{n}_0^{\text{EXP}}[i]$ and $\mathbf{n}_0[i]$ are the experimental and theoretical directors in corresponding bins $i$, respectively (see Supplementary Fig. 4a), and $N_{\text{bin}}$ is the total number of bins that are occupied with sufficient data in the experimental data set (these are the bins that have a director in them). We calculated $\text{MD}(r_{\text{b}}, r_{\text{f}})$ for all possible combinations of integer pairs $(r_{\text{b}}, r_{\text{f}})$ in the ranges $30 \le r_{\text{b}} \le 160mm$ and $r_{\text{b}} \le r_{\text{f}} \le 400mm$ (as visualized in Supplementary Fig. 4b). The lower bound of $r_{\text{b}}$ was taken to be the average body length of an adult *T. clavipes* female[1], while the upper bounds of $r_{\text{b}}$ and $r_{\text{f}}$ were estimated from the experimental director field. The minimal discrepancy was obtained with $r_{\text{f}} = 218mm$ and $r_{\text{b}} = 30mm$ (see Supplementary Fig. 4c,d), and $C = 1.32$ (by Eq. (S5)). In terms of the parameters of the non-rescaled Eq. (S1), this translates to $\beta/\alpha = 94$ and $C = 1.32$.

### Supplementary Note 4: Langevin parameters estimation

Recall the Langevin equations used to simulate the motion of real males (Eqs. (2) and (3) of the main text),

$$\frac{\partial}{\partial t}\mathbf{r}(t) = -\frac{1}{\gamma}\nabla V(\mathbf{r}(t)) + v_{\mathrm{p}}\begin{pmatrix}\cos(\phi(t))\\ \sin(\phi(t))\end{pmatrix} + \sqrt{4D_T}\begin{pmatrix}\xi_x(t)\\ \xi_y(t)\end{pmatrix}, \tag{S15}$$

$$\frac{\partial}{\partial t}\phi(t) = \sqrt{2D_R}\,\xi_\phi(t) \tag{S16}$$

To correctly describe the dynamics of males, we need to estimate the values of $\gamma$, $v_{\mathrm{p}}$, $D_T$, and $D_R$. These values are essentially determined by the interplay between the deterministic nature of the motion ($\gamma$ and $v_{\mathrm{p}}$) and the extent of stochasticity ($D_T$ and $D_R$). This interplay can be evaluated from the directional persistence in male trajectories. We thus compared the autocorrelation of the direction of movement along male trajectories in single-male experiments and simulations. The values of the Langevin parameters were fixed such that these autocorrelation profiles with respect to both time ($\Delta t$) and distance ($\Delta|\mathbf{r}|$) are in good agreement (Supplementary Fig. 5). The autocorrelation was calculated as in [2], with a time window size of $4sec$ for direction calculation. We thereby obtained the values $\gamma = 2/sec$, $v_{\mathrm{p}} = 2\ mm/sec$, $D_T = 5\ mm^2/sec$, and $D_R = 0.05\ rad^2/sec$.

### Supplementary Note 5: Necessity of the medium-range spring-like regime in the male potential

Recall the male effective potential given in Eq. (4) of the main text,

$$V_{\mathrm{male}}(d) = \begin{cases} \frac{\delta}{d} + \frac{k}{2}(d-d_0)^2 + k_0, & d \le d_{\mathrm{s}} \\ \frac{\delta}{d} + \frac{\tilde{k}}{2}(d-\tilde{d}_0)^2 + \tilde{k}_0, & d_{\mathrm{s}} < d \le d_{\mathrm{m}} \\ \frac{\delta}{d}, & d_{\mathrm{m}} < d \end{cases} \tag{S17}$$

To emphasize the necessity of the medium-range spring-like regime ($d_{\mathrm{s}} < d \le d_{\mathrm{m}}$) in reproducing the experimental inter-male velocity profile, we consider a male potential in which this regime is omitted,

$$V_{\mathrm{male}}(d) = \begin{cases} \frac{\delta}{d} + \frac{k}{2}(d-d_0)^2 + k_0, & d \le d_{\mathrm{s}} \\ \frac{\delta}{d}, & d_{\mathrm{s}} < d \end{cases} \tag{S18}$$

The force profiles derived from Eqs. (S17) and (S18) are compared in Supplementary Fig. 6a. We extracted the inter-male velocity profile in two-male simulations (as described in the main text) with the male potential as in Eq. (S18), and compared it to velocity profile extracted from experiments, and to the one extracted from simulations with the male potential as in Eq. (S17) (Supplementary Fig. 6b). The values of all the shared parameters were kept the same for both male potentials. This comparison clearly demonstrates that without the medium-range spring-like regime, the inter-male velocity profile lacks an essential feature, namely, a region of positive inter-male velocity at the respective inter-male distance range.

### Supplementary Note 6: Larger males are faster

In our experiments, we find that the average speed of a male on the web increases with its size (Supplementary Fig. 8). For simplicity, we fitted a line to the speed-size relation,

$$s(w) = \mathrm{m}\left(\frac{w}{w_{\mathrm{ref}}} - 1\right) + \mathrm{n} \tag{S19}$$

where $w_{\mathrm{ref}}$ is a reference male's weight. With $w_{\mathrm{ref}} = 7mg$, we obtained the corresponding fit parameter values $\mathrm{m} = 0.27\ mm/sec$ and $\mathrm{n} = 1.34\ mm/sec$. Rescaling Eq. (S19) by n, such that the average speed of a male with the chosen reference weight is defined as '1', we can write Eq. (S19) as

$$\bar{s}(w) = a\left(\frac{w}{w_{\mathrm{ref}}} - 1\right) + 1 \tag{S20}$$

where $a = \mathrm{m}/\mathrm{n} = 0.2$. Eq. (S20) is equivalent to Eq. (6) of the main text,

$$\eta_i = a\left(\frac{\delta_i}{\delta_{\mathrm{ref}}} - 1\right) + 1 \tag{S21}$$

if we assume that $\delta_i$ is proportional to the weight of male $i$, so that the ratio between the vibration-based repulsion strengths of two males is simply the ratio of their weights. We note that choosing a different reference weight withing the sampled range does not change the value of $a$ substantially. The value $a = 0.2$ was therefore used for the simulations of different sized males.

### References


[1] Leonor Ceballos Meraz, Yann Hénaut, and Mark A Elgar. "Effects of male size and female dispersion on male mate-locating success in Nephila clavipes". In: *Journal of ethology* 30.1 (2012), pp. 93–100.

[2] Roman Gorelik and Alexis Gautreau. "Quantitative and unbiased analysis of directional persistence in cell migration". In: *Nature protocols* 9.8 (2014), p. 1931.
