## Supplementary figures and images for "Dynamics on the web: spiders use physical rules to solve complex tasks in mate search and competition"

### caption for Supplementary Movie 1

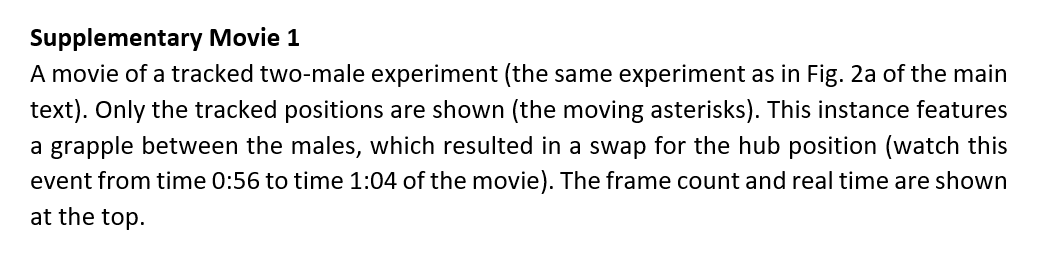
